# Lung function candidate genes in *Drosophila melanogaster*

**DOI:** 10.64898/2026.06.10.731340

**Authors:** Emma Szamek, Zsuzsa Markus, Marina Molina Altamirano, Núria Corbella-Rius, Ian Adcock, Sofia J. Araújo, Ian Sayers, Marios Georgiou

## Abstract

**Rationale:** Chronic obstructive pulmonary disease (COPD) represents a leading cause of global morbidity and mortality. Genome-wide association studies (GWAS) have implicated numerous genetic variants in lung function impairment, yet confidently identifying the underlying genes and pathways, and translating these findings into mechanistic insight, remains a significant challenge.

**Objectives:** To leverage the genetic amenability and high-throughput screening capability of *Drosophila melanogaster* to determine the role of candidate causal genes in epithelial cell homeostasis.

**Methods:** We performed a loss-of-function analysis of 60 prioritised lung function candidate causal genes implicated from GWAS in two distinct epithelia: the dorsal thorax and trachea.

**Results:** We identified 57/60 tested candidate genes that alter at least one aspect of epithelial morphology and behaviour upon knockdown. With a focus on junctional integrity, cell delamination and tissue growth, we identified 11 genes for further study: *Sec6, RpS26, pAbp, Arf102f, Riok1, Sra-1, Inpp5e, CG31759, ssh, eIF6 and Rtf1*. Further characterisation found a significant reduction in junctional E-Cadherin levels following *Arf102F*, *Rtf1*, *RioK1* and *Sra-1* knockdown. Following a secondary screen in the *Drosophila* tracheal system for priority candidates, *Sec6* and *RpS26* were associated with significant airway defects and a reduction in larval body size. 8/11 priority genes exhibited differential lung gene expression between controls and patients with COPD.

**Conclusions:** These data demonstrate the amenability of *Drosophila* melanogaster to perform *in vivo* functional analyses of candidate causal genes at scale. Initial findings implicate several genes in epithelial homeostasis and integrity, providing new mechanistic understanding and potential therapeutic targets for COPD.

**Graphical Abstract:** 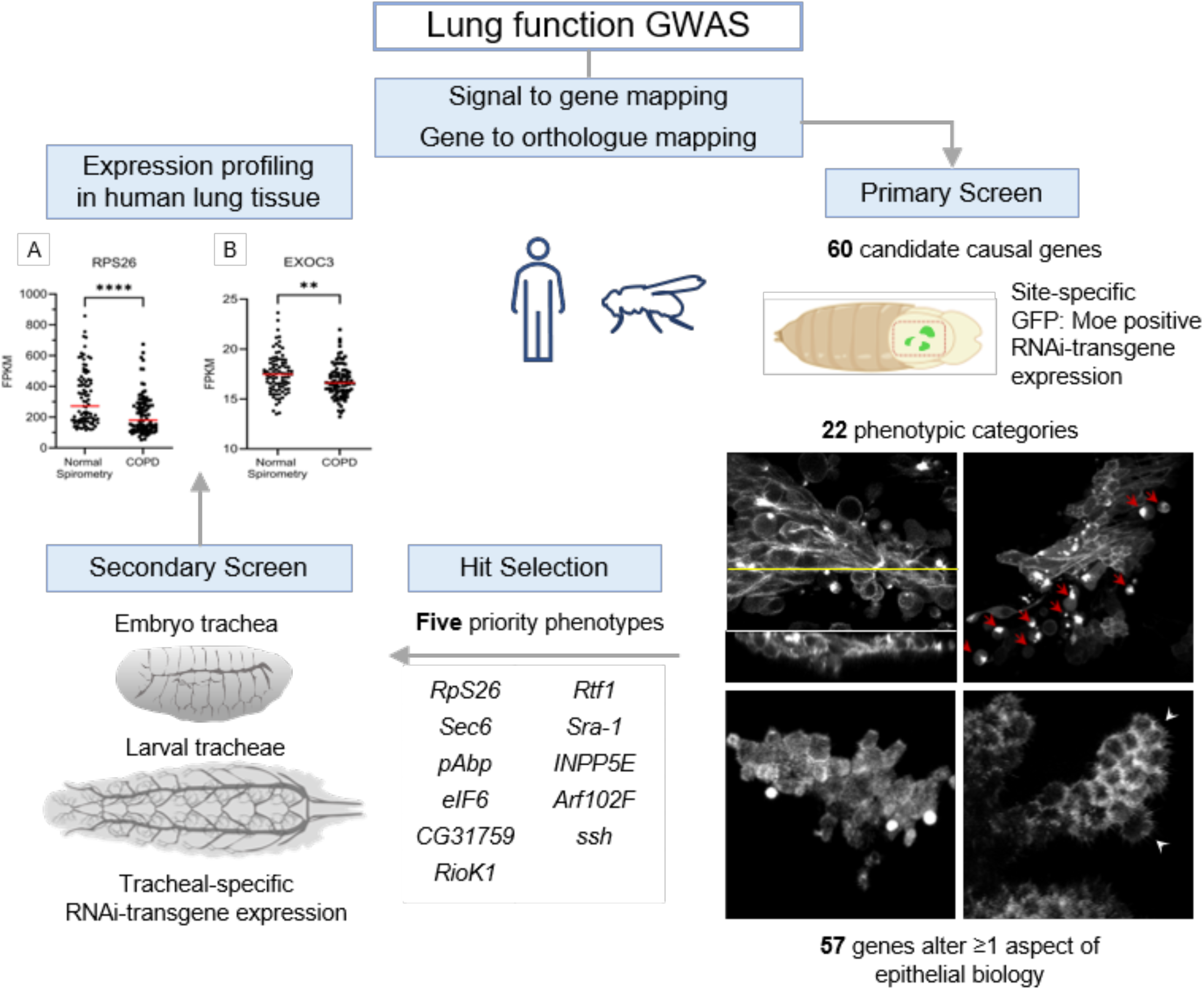

## Introduction

Chronic obstructive pulmonary disease (COPD) is a heterogeneous lung condition marked by a progressive irreversible airflow obstruction (1). The airway epithelium serves as a frontline defence against inhaled insults, fulfils multiple functions to maintain pulmonary homeostasis, and coordinates intercellular signalling within the airway microenvironment. In chronic respiratory diseases such as asthma, idiopathic pulmonary fibrosis (IPF) and COPD, the structure and biology of the airway epithelium are modified, including consequences for barrier function (2,3).

COPD susceptibility and heterogeneity are attributed to dynamic interactions between genetic and environmental factors, such as cigarette smoking and biomass smoke exposure, across the life course. Genome-wide association studies have implicated a large number of genetic variants influencing risk to develop COPD, lung function impairment, and related phenotypes (4–6). Given the close association between lung function decline and COPD onset (7,8), the identification of genetic factors, candidate causal genes, and pathways that alter lung function through changes in epithelial biology has the potential to yield new mechanistic insights and identify novel therapeutic targets.

Previously, functional genetic studies have favoured the use of mammalian models, particularly transgenic mice, which have a high associated cost, time burden, and impact on animal welfare. The fruit fly *Drosophila melanogaster* has proven to be a powerful, cost-effective model for studying human disease, including lung research (9). Its high degree of genetic conservation, short lifespan, and advanced genetic toolkit enable rapid, multigenerational studies and make it an ideal *in vivo* screening tool. Importantly, it has comparatively simpler epithelial tissues with which to study orthologous genes in the absence of adaptive immunity (10–12).

In this study, we conducted a detailed systematic loss-of-function (LOF) screen for prioritised candidate causal genes associated with lung function impairment by integrating advanced *Drosophila* genetic techniques with transgenic RNAi technology. We employed a two-stage screening protocol, using two distinct *Drosophila* epithelia: an initial screen in the dorsal thorax epithelium, followed by a secondary screen in the fly’s tracheal system. The dorsal thorax is an epithelial sheet that is highly accessible and allows for the high-resolution real-time imaging of epithelial shape and behaviour (13–15). The tracheal system consists of a network of epithelial tubules that functions as the fly’s respiratory organ.

We demonstrate that the fruit fly is an effective *in vivo* screening tool and identify multiple genes with significant roles in regulating epithelial stability and structural organisation, offering a platform for translating GWAS findings into mechanistic and therapeutic insight.

## Methods

Additional details of methods are provided in the online supplement.

### Prioritisation of candidate causal genes for functional study

At the time of analysis, the largest GWAS meta-analysis of lung function impairment utilised 19,819,130 SNPs genotyped in UK Biobank and the SpiroMeta consortium comprising over 400,102 individuals of European ancestry (5). In this study, 279 signals were significantly associated with four quantitative lung function traits: FEV_1_, FVC, FEV_1_/FVC and PEF. For the current study, we used an integrative *in silico* analysis pipeline to prioritise candidate genes for investigation *in vivo* from these genetic association data (Supplementary Methods).

### *Drosophila* screening of 60 prioritised candidate genes

We completed an *in vivo* knockdown genetic screen of the 60 candidate causal genes in the fly allowing us to: (1) generate a patch of tissue on the dorsal thorax that overexpresses an RNAi transgene to attenuate the expression of the gene of interest, surrounded by wild-type (WT) tissue; (2) specifically label the knockdown tissue with GFP:Moe (the actin-binding domain of moesin fused to GFP) (16) as shown in Figure 1.

**Figure 1:**
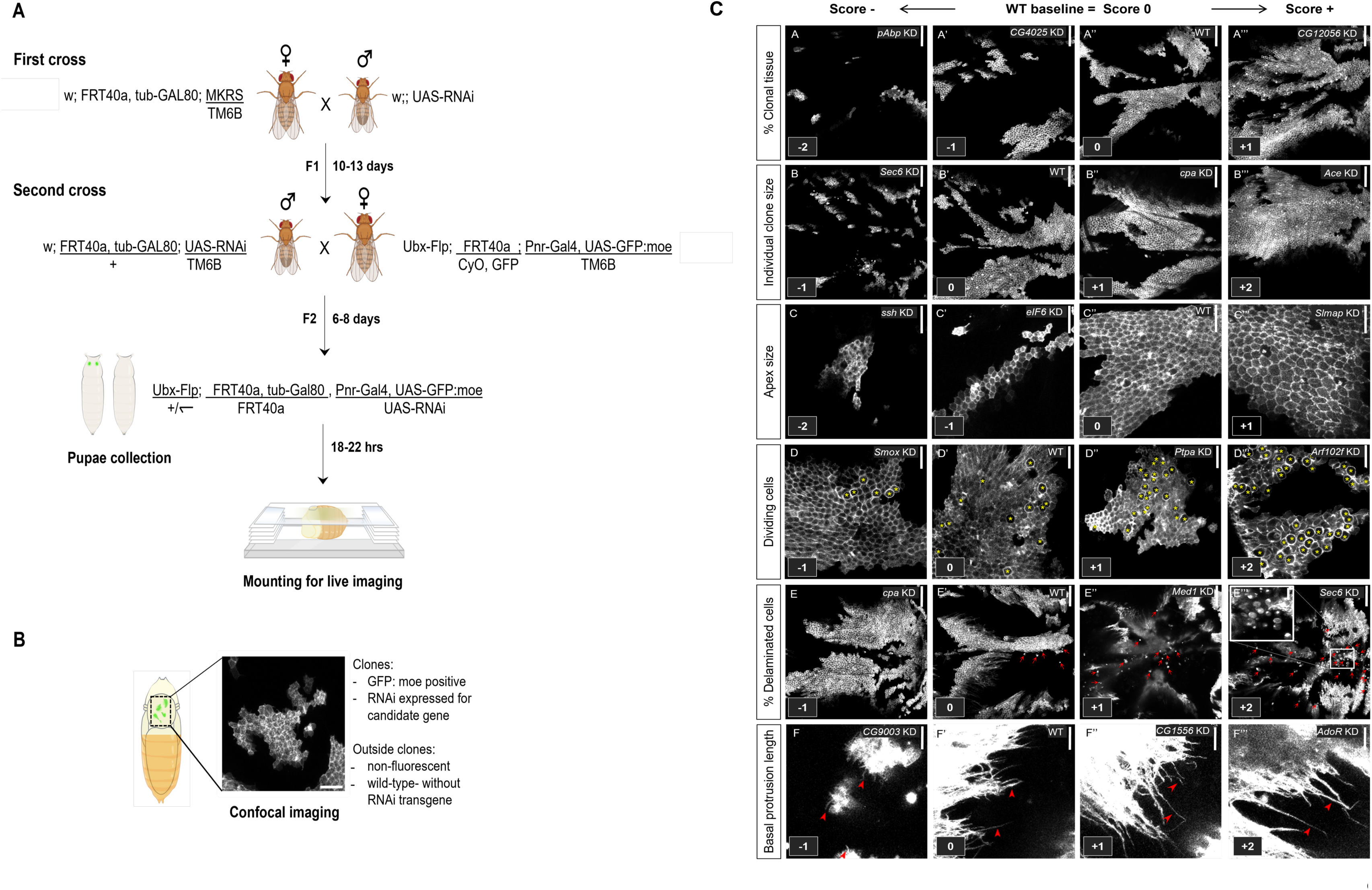
Screening approach for the primary screen. (A) First-generation crosses used transgenic females carrying an FRT site and tubGal80 with UAS-RNAi males. F1 progeny of the appropriate genotype, specifically MKRS-negative males were crossed with females containing a complementary FRT site, Flp recombinase, Pnr-Gal4 and UAS-GFP:moe. F2 pupae were collected at 0hr APF and selected for a non-tubby shape (i.e., lacking the TM6b balancer) and GFP-positive clones in the notum (non-fluorescent pupae were also collected). See (16) for a detailed explanation of the genetic constructs and balancers used. (B) Selected pupae were aged at 29°C and mounted for live imaging from 18-21 hr APF. The notum is enclosed (dashed black rectangle) and a representative confocal image shows GFP:moe-positive RNAi-expressing clones adjacent to non-fluorescent wild-type cells. Scale bar: 20 μm(C) Our semi-quantitative scoring system applied to several phenotypes. Representative images of WT background phenotypes scored zero; positive and negative scores indicate relative increase and decrease in phenotype, respectively. Genotypes are indicated at the top right corner of each panel. Yellow asterisk pinpoints dividing cells; white box highlights a magnified view of delaminating cells. Red arrows, delaminating cells; red arrowheads, basal protrusions. Scale bar: 100μm (A-A’’’, B-B’’’, E-E’’’); 20μm (C-C’’’, D-D’’’, F-F’’’).

### Dissections and live imaging

Pupal aging, mounting, and dissections were performed as previously described (13). Imaging of live pupae and fixed samples was performed with a Zeiss LSM880 inverted confocal microscope equipped with a x40/1.30 NA oil Ph3 M27 objective; z-series were acquired with 1μm z-sectioning for live pupae and 0.5μm for fixed samples. Imaging of immunostained embryos and larvae was carried out using a Zeiss AxioPhot fluorescence microscope and the Leica DMLB fluorescence microscope respectively. All images were processed and quantified in Fiji ImageJ (3.1.0) software (17).

### Immunostaining

We used the following primary antibodies at the indicated dilutions for this study: rat anti-E-Cad [1:100, DSHB (DCAD2)], mouse anti-Gasp [1:5 DSHB (2A12)], an HRP-conjugated secondary antibody and a secondary antibody Alexa Fluor® 647 IgG goat anti-rat [1:300, Abcam].

### Semi-quantitative scoring of phenotypes

Semi-quantitative scoring was an adaptation of the method used in a previous large-scale screen (13). For each phenotype, a numerical score indicates the degree of similarity or difference between the subject animal and a WT control animal, assigned a score of zero. Any observed change was denoted by a positive or negative score scaled according to the degree of enhancement or reduction of the phenotype respectively. For binary categories (yes/no) a given score can signify the presence (1) or absence (0) of a phenotype e.g. closure defects, apical defects, basal bundles. For a full description of the scoring system see Supplementary Methods.

### Expression profiling of 11 prioritised candidate genes in human lung

To determine whether the expression levels of our prioritised candidate genes are altered in the lungs of patients with COPD, we utilised the Gene Expression Omnibus (GEO) dataset GSE57148, containing gene expression data for lung tissue taken from patients with normal spirometry (n=91) and patients with a diagnosis of COPD (n=98) (18).

### Statistical analysis

Microsoft Excel and PRISM (GraphPad) software were used to perform calculations, generate graphs, and calculate statistical significance, with Student’s *t*-test, Mann–Whitney U test, and Spearman’s Correlation test. ns *P* ≥ 0.05, \**P*< 0.05, \*\**P*< 0.01, \*\*\**P*< 0.001,\*\*\*\**P*< 0.0001; *r_s_* [±0.7, ±1.0] strong, *r_s_* [±0.5, ±0.7] moderate, *r_s_* [±0.3, ±0.5] weak, *r_s_* [0, ±0.3] negligible correlation.

## Results

### In silico analyses identify 60 candidate causal genes for study *in vivo*

In our GWAS signal-to-gene framework we focused on the 70 signals that had the greatest confidence for variant-to-gene mapping in the original GWAS (5), from which we identified 187 potential candidate causal genes that were supported by at least two independent lines of evidence (Table S1-S8). Cross-species mapping identified 85 high-confidence *Drosophila* orthologues, which displayed consistent expression throughout embryonic, larval, and pupal development, particularly within epithelial precursors of the dorsal thorax (Table S9, S10). Subsequently, we obtained 78 UAS-RNAi lines corresponding to 60 human candidate causal genes in total for loss-of-function analysis (Table S11).

### Systematic loss-of-function analysis in the pupal notum identifies 57/60 genes which alter epithelial characteristics

We applied an unbiased approach to identify genes that increase or decrease specific aspects of epithelial shape, dynamics and behaviour in our system (Figure 1). We generated a database consisting of 22 phenotypic categories (Table S13), where each animal with RNAi KD clones was scored relative to animals with WT clones. Each category describes an aspect of epithelial cell behaviour. Categories include clone size and shape, number of dividing cells, number of delaminating cells, apex size, junction defects, cytoskeletal defects, multilayering, etc. The scoring system we employed reflected the fact that gene KD could either positively or negatively affect specific aspects of cellular behaviour (Figure 1C; Table S13). We calculated the mean score for each of the 78 RNAi lines across each of the 22 distinct phenotypic categories (Figure 2, Table S13). Using these averages, we determined the distribution of scores for all categories. Genes with a mean score ±2 SD above or below the WT baseline were selected as genes of interest, yielding 57 unique genes of interest (Tables S13-37).

**Figure 2:**
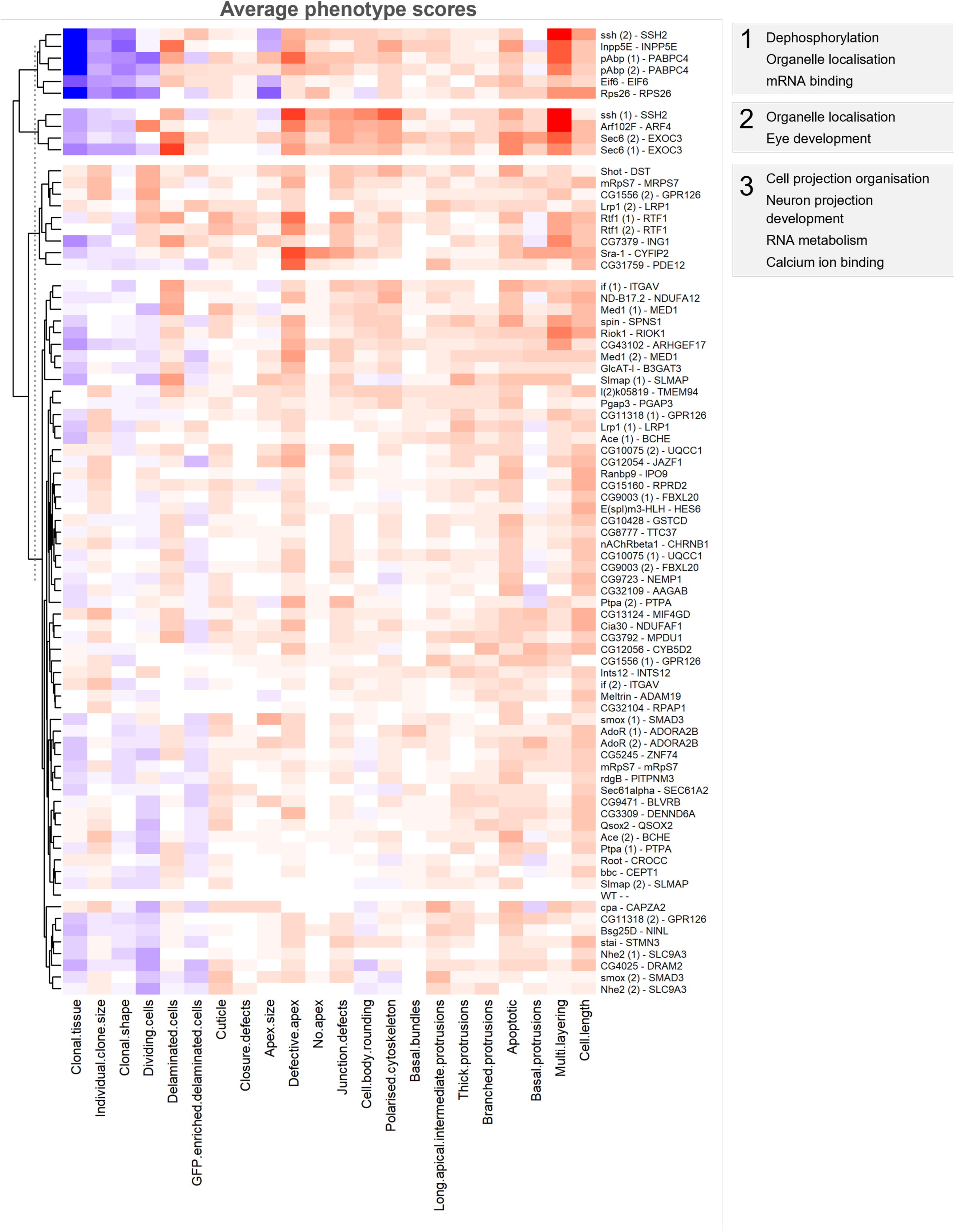
A heatmap representation of average phenotype scores for all 78 RNAi lines. Each row represents an RNAi line which is labelled as *Drosophila* orthologue - human gene; each column represents a phenotype category. Genes targeted by two independent RNAi lines are denoted (1) and (2). Rows are clustered by hierarchical analysis of phenotype score direction and magnitude. Map colours represent row-scaled average scores: blue indicates the lowest score, light blue indicates an intermediate score, and red indicates the highest score. Each cluster was analysed with regard to their biological and molecular function by GO enrichment analysis. The top significant GO categories after multiple testing correction (Benjamini–Hochberg FDR < 0.05) are displayed for each cluster.

To verify that our semi-quantitative scoring system reflected meaningful results that represented real phenotypic differences, RNAi lines were selected at random and subjected to full quantitative analysis across multiple phenotypes considered the most amenable to in-depth quantification: average percentage of clonal tissue, invading cells, dividing cells, and the length of basal protrusions. A strong positive correlation was observed for all categories measured (0.92–0.98, Spearman correlation test) (Figure S1). We also considered whether genes targeted by two independent RNAi lines produced similar phenotypes (Figure S1). We compared average scores across 22 phenotypic categories for each pair of RNAi lines and determined that 9/15 genes targeted by two independent RNAi lines demonstrated statistically similar phenotypes. Thus, we have a relatively high degree of confidence that the observed phenotypes are the result of depleted gene expression rather than due to off-target effects.

To establish whether gene knockdowns gave rise to similar phenotypes, unsupervised hierarchical clustering was performed on all RNAi lines studied across all phenotypes. This resulted in the identification of four distinct clusters shown in Figure 2. In parallel, we explored the potential biological relevance of genes within each cluster by GO-term enrichment analysis, which identified significantly enriched terms after multiple testing correction (Table S12).

### Gene Prioritisation for in-depth analyses

Five phenotypic categories were identified as most relevant to COPD pathobiology: apical defects, junctional defects, cell delamination, multilayering, and reduced clonal tissue size. We also interrogated penetrance to help prioritise genes for further study (see Supplementary Methods). As shown in Figure 3, utilising this approach, we identified 11 candidates of interest which strongly and reproducibly disrupt the dorsal thorax epithelium: *Arf102F, CG31759, Inpp5e, pAbp, RioK1, RpS26, Rtf1, Sec6, Sra-1, ssh and eIF6*.

**Figure 3:**
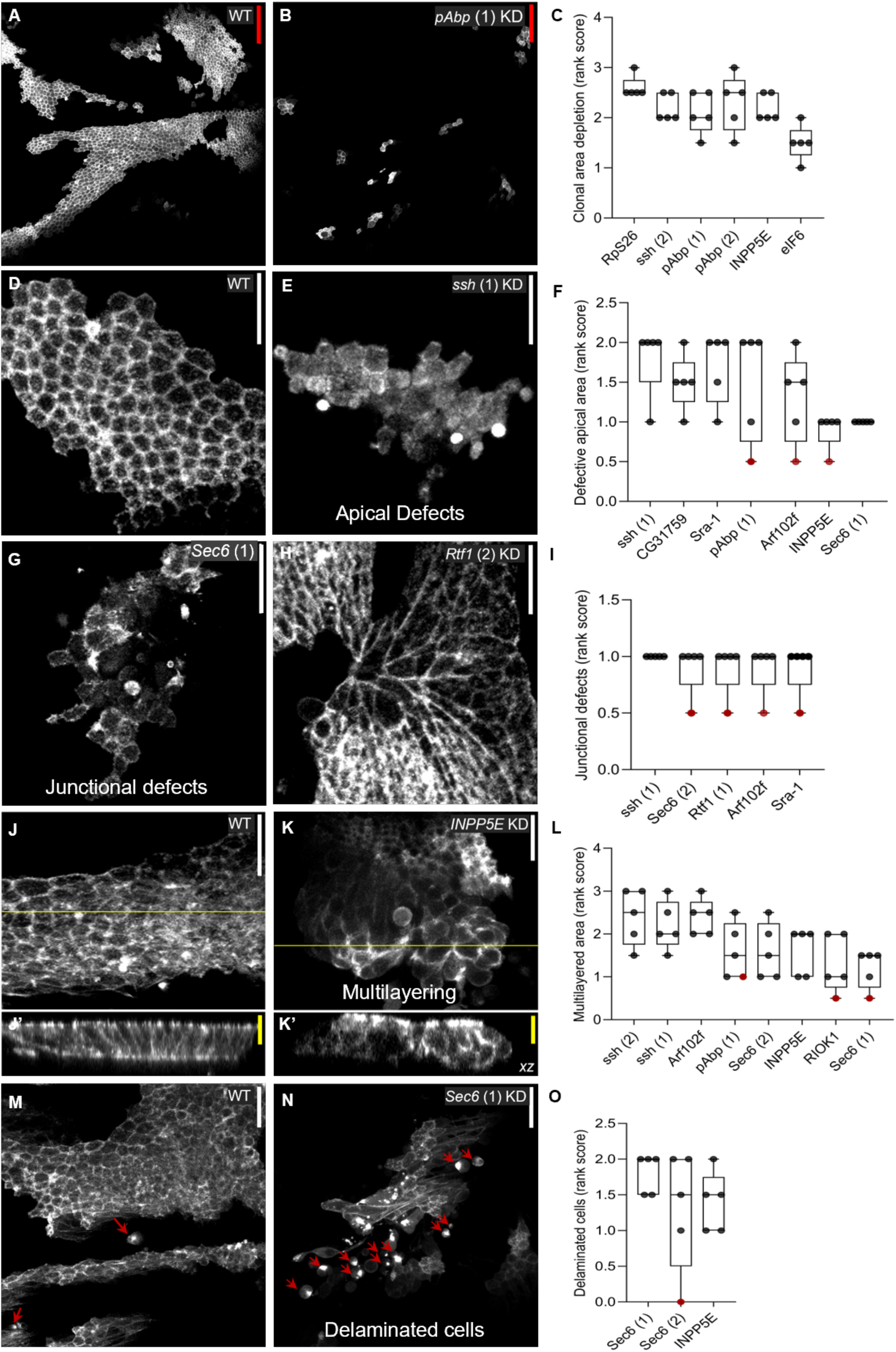
11 high-priority candidate genes identified in five key categories disrupting epithelial homeostasis. (A-B, D-E, H, J-K, M-N) GFP:moe-labelled genetic clones in the dorsal thorax epithelium of living fly pupae. Yellow line: cross section of xy plane; red arrow: delaminated cell. Yellow scale bar: 10 μm; white scale bar: 20 μm; red scale bar: 50 μm. (C, F, I, L, O) Ranked scores in each selected phenotypic category. Shown are the genes prioritised based on average scores ± 2 SD from the wildtype average and concordance between scorers for at least 4 of 5 animals screened per line. Each dot represents 5 animals. Black dots: concordant scores; red dots: non-concordant scores between scorers.

### Depletion of *Arf102F*, *Rtf1*, *RioK1* and *Sra-1* lowers junctional E-Cadherin levels

To further delineate the molecular mechanisms by which loss of candidate gene expression influences cell-cell adhesion, we analysed *Drosophila* E-cadherin localisation in the notum. We observed abnormal junctional morphology in *Inpp5e*, *Arf102F*, *RpS26*, *Sec6,* and *Sra-1* knockdown lines (Figure 4). On quantitative analysis, we found that the knockdown of *Riok1*, *Sra-1*, *Arf102F* and *Rtf1* resulted in significantly lower levels of E-cadherin at cell junctions (Figure 4). For two secondary lines, *pAbp* (#28821) and *ssh* (#38948), we were unable to quantify fluorescence due to the very low percentage of clonal tissue (not shown). Although *Sec6* knockdown did not appear to significantly alter E-cadherin levels, mis-localisation was clearly observed for both independently targeting lines.

**Figure 4:**
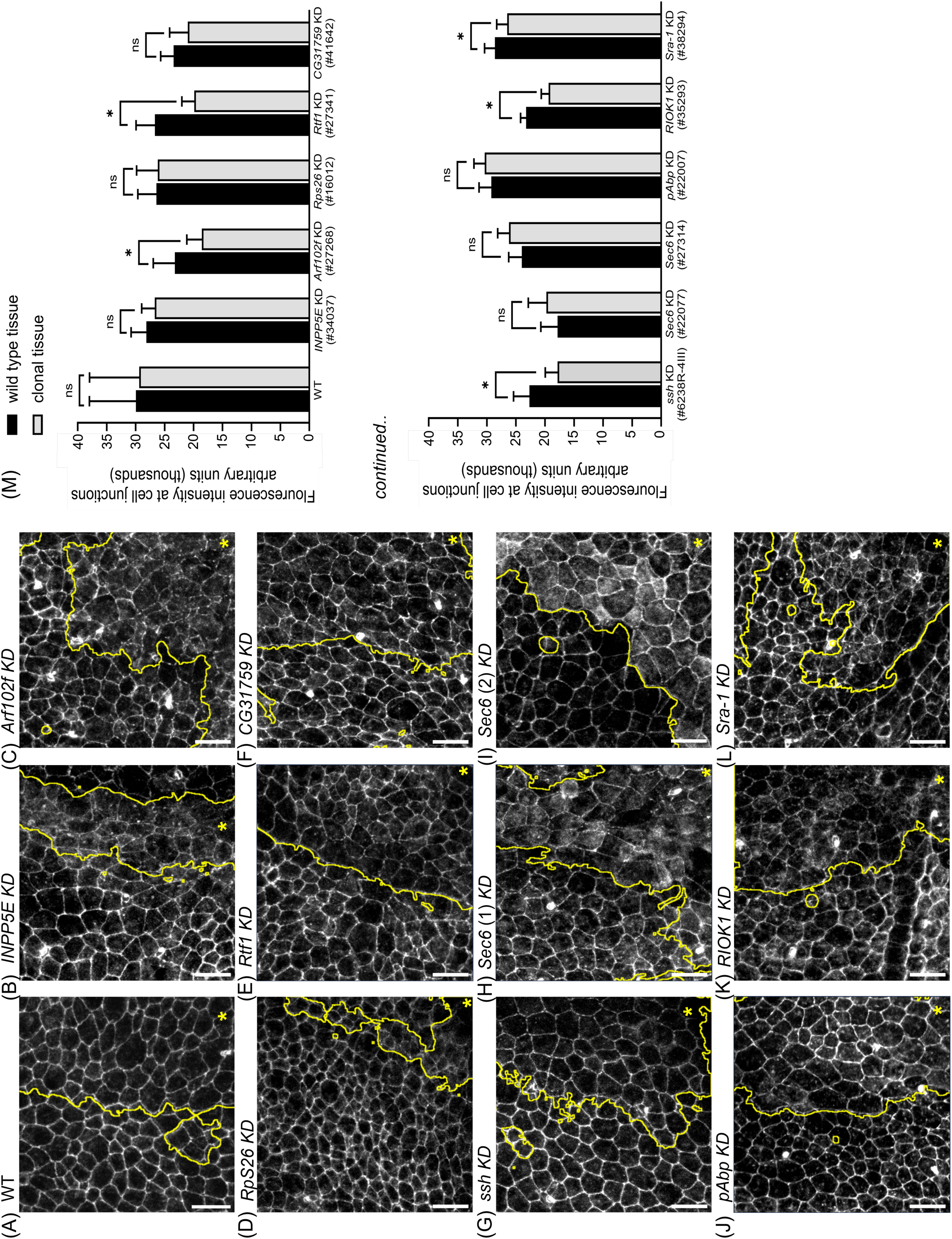
Depletion of Arf102F, Rtf1, RIOK1 and Sra-1 lowers junctional E-Cadherin levels. Junctional staining of ten screen hits indicates loss of E-cadherin and possible mis-localisation. (A-N) *Drosophila* pupal nota containing positively marked clones for Rtf1 KD, ssh KD, Sra-1 KD, pAbp KD, RIOK1 KD, RpS26 KD, CG31759 KD, INPP5E KD, Sec6 KD and Arf102F KD (enclosed on the right-hand side of the demarcated border and marked with an asterisk) were stained for E-cadherin. (M) Following quantification of junctional fluorescence, a significant disruption in junction protein localisation and junction integrity was found in multiple knockdown lines (n ≥ 65 junctions from a minimum of six animals for each genotype). Scale bars = 10 µm. Error bars represent intensity average ± s.e.m. Student’s t-test was performed to determine statistical significance. \**P* < 0.05.

### *RpS26*, *Sec6* and *pAbp* knockdown results in significant larval tracheal abnormalities

To study the effect of candidate gene knockdown on airway integrity, we used the GAL4-UAS system to induce respiratory system specific gene downregulation (19). For these analyses, data for 8/11 were available with 3/11 not available. From embryonic stages 13 to 17, no significant tracheal phenotypes in *Sec6* and *RpS26* knockdown embryos were observed (Figure S2). We then proceeded to assess overall fitness and tracheal morphology at later developmental stages. For *pAbp*, *Rtf1*, *Sec6 and RpS26* knockdown larvae, animals proceed through embryonic development but die as growth-arrested third instar larvae. In time-lapse recordings through instars 1 - 3 (Figure S3), tracheal alterations were observed in *RpS26* (#16012), *Sec6* (#22077, #27314) and *pAbp* (#28821). By the third larval instar, knockdown of *RpS26*, *Sec6* and *pAbp* (Figure S3) gave rise to tracheal dorsal trunk defects characterised by thinner tubular width. In addition, a discontinuous tubular structure was observed in the dorsal trunks of *RpS26* (#16012) and *Sec6* (#22077) knockdown larvae (Figure 5). This phenotype was highly penetrant in both lines (>70% larvae, n =30) and is consistent with a loss of airway integrity.

**Figure 5:**
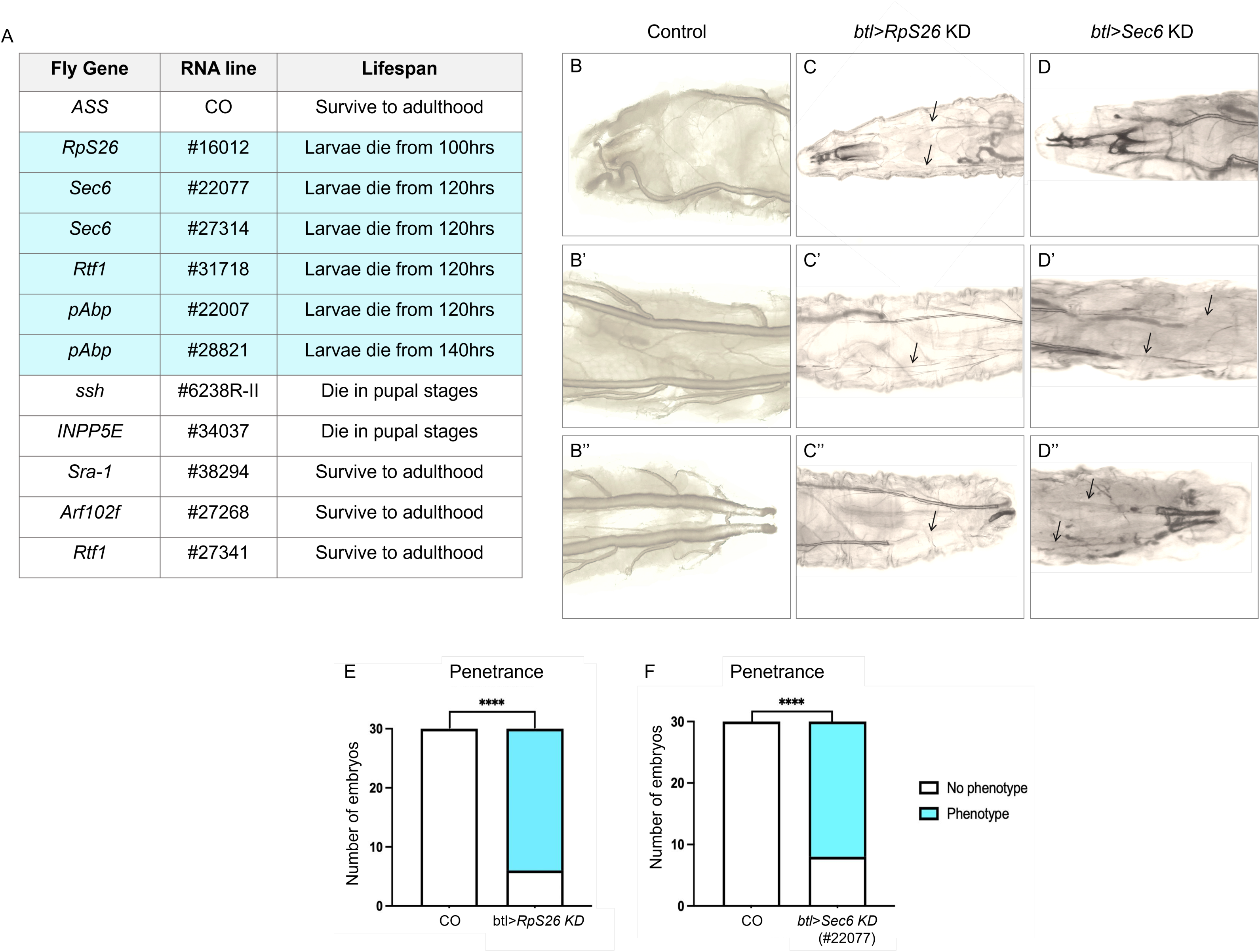
Significant larval instar 3 (L3) tracheal defects were observed in RpS26 and Sec6 knockdown lines. (A) A summary of the lifespan for each btl>RNAi line including larval (green) and pupal (brown) phases. (B-B” – D-D’’) Relative to the negative control (#47097) (B-B”), btl>RpS26 knockdown (#16012) L3 larvae (C-C”) and btl>Sec6 (#22077) knockdown L3 larvae (D-D’’) present airway narrowing and a discontinuous lumen that lacks air in multiple segments. (E) Tracheal phenotypes for both gene knockdowns are highly penetrant. Dorsal view. n (number of larvae) = 30. \*\*\*\**P* < 0.0001.

### Transcriptomic analysis of prioritised candidate genes in human lung tissue from patients with COPD and control subjects

To begin the translation of the 11 prioritised candidate genes, we utilized RNA-seq dataset GSE57148 available on the Gene Expression Omnibus (GEO) and expression profiled these genes in human lung tissue from patients with COPD and control subjects. Human orthologues are referred to using standard HGNC nomenclature. Eight out of ten priority genes exhibited differential expression between patient groups. As summarised in Figure 6, *INPP5E*, *RPS26*, and *EXOC3* expression was significantly lower in COPD lung tissue compared with healthy controls. By comparison, *CYFIP2*, *SSH2*, *RTF1*, *PDE12* and *PABPC4* expression was significantly higher in COPD lung tissue relative to healthy controls. The largest differences between groups were observed for *RPS26* (0.66-fold decrease, *P* < 0.0001), *SSH2* (1.39-fold increase, *P* = 0.016) and *PDE12* (1.65-fold increase, *P* < 0.0001). The concordance between functional phenotypes identified in *Drosophila* and dysregulated gene expression in human COPD lung tissue supports the translatability of findings from this model and strengthens the candidacy of these genes as contributors to COPD pathobiology.

**Figure 6:**
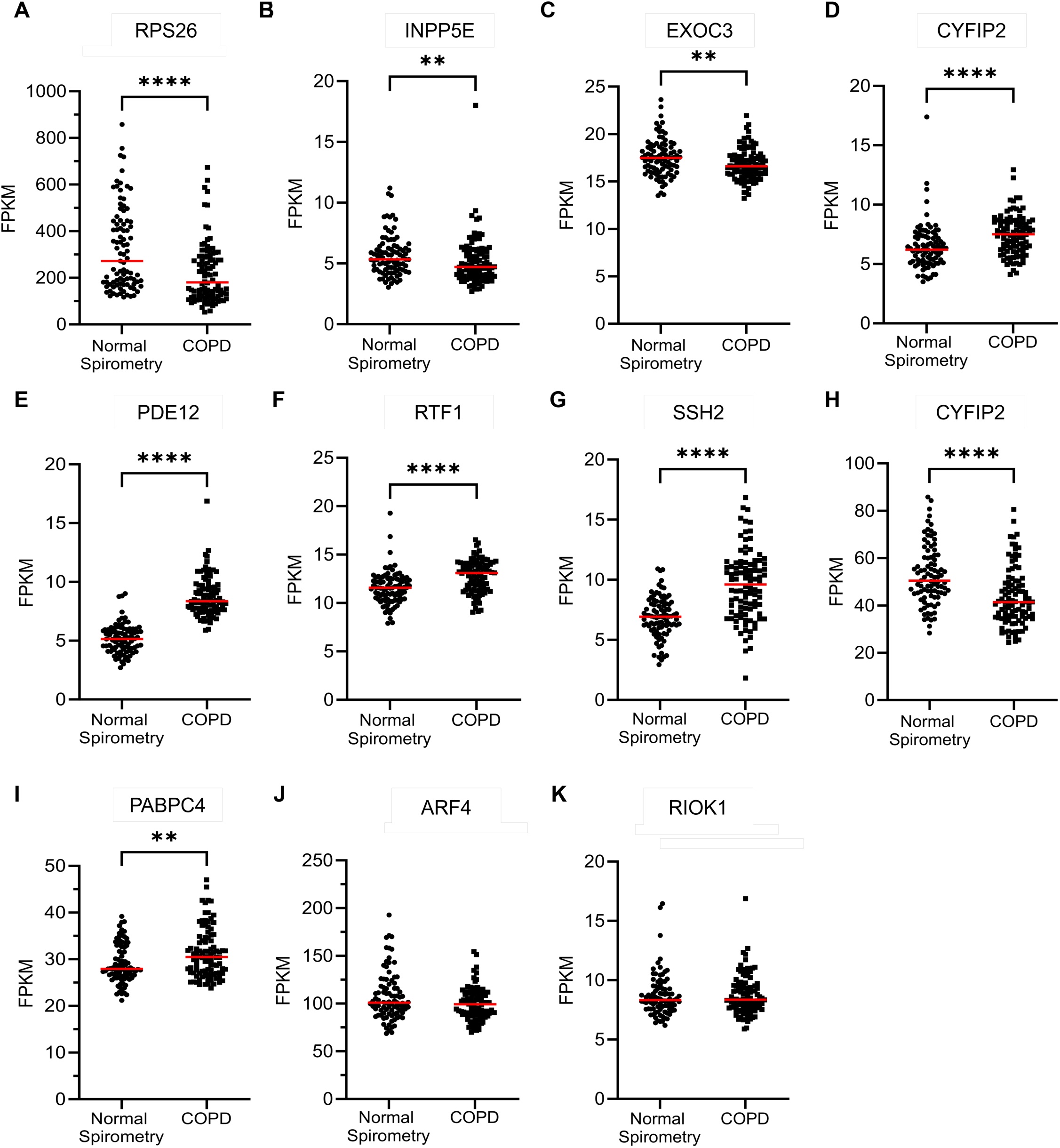
Expression profiling of the top 11 candidate genes in human lung tissue from patients with COPD (n = 98) and subjects with normal spirometry (n = 91). Transcript abundance (FPKM, fragments per kilobase per million) is shown for (A) *RPS26*, (B) *INPP5E*, (C) *EXOC3*, (D) *CYFIP2*, (E) *PDE12*, (F) *RTF1*, (G) *SSH2*, (H) *EIF6*, (I) *PABPC4*, (J) *ARF4*, (K) *ARHGEF17*, and (L) *RIOK1*. Red horizontal lines indicate the median. Expression values were taken from the dataset GSE157148 and a Mann-Whitney test with two-stage linear step-up procedure of Benjamini, Krieger and Yekutieli used to control the FDR at 5% was performed. \**P* < 0.05, \*\**P* < 0.01, \*\*\**P* < 0.001, \*\*\*\**P* < 0.0001

## Discussion

There has been significant progress in identifying genetic variants associated with lung function impairment and COPD, however, translation to genes, pathways and new understanding has been a challenge. In this study we set out to leverage the genetic tractability of *Drosophila melanogaster* in combination with RNAi-mediated transgenics to allow us to interrogate candidate causal genes on scale for impact on epithelial cell biology. We illustrate the utility of this platform and strikingly, 57/60 prioritised candidate genes alter at least one epithelial characteristic. Focusing on a subset of these genes that influenced five core epithelial functions we considered of greater relevance to COPD, we show significant effects on junctional morphology and E-cadherin driven by loss of *Rtf1*, *Arf102f* (*ARF4*), *RioK1* (*RIOK1*) and *Sra-1* (*CYFIP*). To translate further to the airway, we demonstrate significant effects on the development of the fly trachea (*RpS26* and *Sec6* (*EXOC3*)) and show that the majority of prioritised genes are differentially expressed in lung of patients with COPD. Taken together, our study demonstrates the significant utility of this *in vivo* platform and identifies new candidate causal genes for future study.

Using the most recent GWAS of lung function impairment at the time (5), we systematically identified 60 candidate causal genes for *in vivo* study. In the initial dorsal epithelial loss-of-function screen, 57 of these candidates impacted epithelial characteristics, highlighting the potential enrichment of genes related to epithelial function in GWAS of lung function, as previously suggested (4,20), a finding that warrants further investigation. We focused on a subset of gene hits for junctional defects, apical defects, cell delamination, multilayering, and reduced clonal tissue size as we considered these phenotypic measures to be particularly relevant to COPD.

Our results indicate that targeting Sra-1, Arf102f, and Rtf1 led to reduced junctional E-cadherin levels. The loss of epithelial integrity, increased permeability, and disrupted expression of apical and tight junctional proteins that underlie junctional abnormalities are well documented in airway epithelial cells derived from COPD patients (2,21,22). Junctional stability is critically dependent on the interaction between cadherins and the actin cytoskeleton. *CYFIP2,* the closest human orthologue of *Sra-1*, is a component of the WAVE regulatory complex (WRC) that promotes F-actin polymerization (23). *CYFIP* knockdown in mammary epithelial cell lines has been linked to aberrant E-cadherin distribution (24). Direct orthologue *Rtf1*, a subunit of the Polymerase-Associated Factor 1 (Paf1) complex, functions as a key regulator of elongation and co-transcriptional histone modification (25,26). Interestingly, loss of Rtf1 activity in cardiomyocytes was associated with fibrosis and defective junctions, with strong evidence of diminished N-cadherin localisation (27), mechanisms known to be important in lung homeostasis. The *ARF4* gene, a close orthologue of *Arf102F,* is a GTPase implicated in vesicular transport, including the sorting and transport of ciliary proteins to the primary cilium (28,29). Notably, *ARF4* has been implicated in the maintenance of intestinal epithelial permeability through the regulation of tight junction proteins such as claudin-4 (30). Although E-cadherin was depleted in the dorsal thorax epithelium following knockdown, these lines remained viable through to adulthood in the secondary screen. This indicates that reduced junctional E-cadherin alone may be insufficient to compromise organismal viability, although further experiments in the fly airway are required.

In the tracheal analyses, we observed in the third instar of *RpS26 (RPS26), Sec6* (*EXOC3*), and *pAbp* (*PABPC4*) knockdown larvae a disruption to the developmental program of tracheal expansion in diameter and length of the tubes. We hypothesise that these defects result in a severe undersupply of oxygen to surrounding tissues which may limit larval growth and confer lethality. The *Sec6* gene encodes a key component of the exocyst complex that mediates the docking of exocytic vesicles with fusion sites on the plasma membrane (31). During embryonic development in *Drosophila*, *Sec6* and *Sec8* localize to the cell periphery of the elongating terminal tracheal cell, with preferential accumulation at the cell tip (32). As terminal branches ramify extensively during the larval stage, this may explain the low penetrance observed in embryos. The larval tracheal defects observed in *Sec6* knockdown lines are consistent with previous findings which determine that the exocyst complex is essential for dorsal trunk elongation and the maintenance of cell size and branch complexity (33,34). Interestingly, in human lung tissue, we demonstrated reduced EXOC3 expression in human lung tissue; decreased expression of *EXOC3* has been shown to reduce gas exchange in the lungs (35). The *RpS26* gene, the direct *Drosophila* orthologue of human RPS26, encodes ribosomal protein S26 and plays a key role in final steps of the maturation of 40S subunits (36). Several RPs perform important extra-ribosomal functions, influencing processes such as cell differentiation, DNA repair and apoptosis (37). In mosaic *Drosophila* tissues, heterozygous mutant clones were found to be actively eliminated in mosaic tissues through ‘cell competition’ in favour of WT populations (38), suggesting that global *RpS26* loss may impair cellular fitness. We demonstrated that *RPS26* gene expression was reduced in the lungs of patients with COPD, however, we have previously reported elevated RPS26 mRNA in bronchial epithelial cells from patients with asthma (39) suggesting disease-specific and context-dependent regulation that warrants further investigation. Comparatively, the *pAbp* gene encodes a poly(A) binding protein, which plays an important role in promoting gene expression by enhancing translation and mRNA stability (40,41). Although the biological consequences of *pAbp* depletion in *Drosophila* tissues are not widely reported, in the prothoracic gland it has been linked to first-instar larval arrest and decreased expression of ecdysteroidogenic genes, that are indispensable for molting and metamorphosis (42). In humans, the homologue *PABPC4* is involved in mitochondrial transport and has been identified as a candidate gene for lung function impairment using multiomics approaches (43) and loss of *PABPC4* has been linked to development of emphysema (44). While these in vivo findings highlight potential mechanisms underpinning the role of these genes in COPD, transcriptomic analysis revealed differential expression of 8 of the 11 prioritised genes in patient lung tissue. These data provide additional support for the involvement of these genes in COPD pathobiology, and highlight the need for further work to determine whether their dysregulated expression is a driver or consequence of disease.

Our study has several strengths: (1) the systematic screening of 60 prioritised candidate causal genes across 22 phenotypic categories represents, to our knowledge, one of the largest in vivo functional genomic screens of GWAS-implicated genes conducted in Drosophila, demonstrating the scalability and efficiency of this platform; (2) the use of an in vivo approach to further our understanding of candidate genes provided significant translational and biological insight at scale; (3) the genes identified provide new insight into how alterations in the epithelium may contribute to COPD pathobiology; (4) most notably, the majority of prioritised genes are also differentially expressed in patient lung tissue, supporting the clinical relevance of the findings.

However, our study does have limitations, most notable, our focus on the functional consequences of downregulating target genes through loss-of-function approaches. While informative, this strategy does not account for the potential effects of overexpression or gain-of-function, particularly for candidate causal genes that exhibited upregulation in expression datasets. While *Drosophila* is advantageous for the study of epithelial remodelling, there are limitations to its applicability to human studies, particularly the lack of an epithelial-inflammatory interface known to be important in human respiratory conditions.

In conclusion, we report what is, to our knowledge, the first systematic use of a *Drosophila* epithelial loss-of-function platform to provide mechanistic insight into candidate causal genes identified in genetic studies of lung function impairment. We identify several candidate causal genes that impact epithelial function, and junctional integrity in particular, that warrant further study and potentially provide opportunities for therapeutic targeting. The concordance between functional phenotypes identified in *Drosophila* and differential gene expression observed in the lung tissue of patients with COPD provides transcriptomic support of our screen and strengthens confidence in the biological relevance of the identified candidates. More broadly, this study demonstrates that functional genomics in *Drosophila* represents a tractable and scalable approach to bridge the gap between GWAS discoveries and disease mechanism, offering a powerful complement to mammalian models in the post-GWAS era.

## Author contributions

I.S. and M.G. conceived the project; M.G., I.S., E.S. and S.J.A. were involved in methodology; E.S., N.C.R., M.M.A. and Z.M. were involved in investigation; E.S., N.C.R., M.M.A. and Z.M. were involved in formal analysis; E.S. was involved in writing - original draft; I.S., M.G. and S.J.A. were involved in writing - reviewing and editing; M.G. and I.S. were involved in funding acquisition; Z.M. was involved in resources; M.G. and I.S. were involved in project supervision.

## Acknowledgements

We would like to thank the fly community for their generosity with reagents, particularly the Bloomington, VDRC and NIG stock centres. We also thank the School of Life Sciences Imaging (SLIM) team for their technical support with confocal microscopy.

## Data availability

All data available on request.

## References

1. GOLD Report. GLOBAL STRATEGY FOR PREVENTION, DIAGNOSIS AND MANAGEMENT OF COPD: 2024 Report [Internet]. 2024. Available from: https://goldcopd.org/2024-gold-report/

2. Heijink IH, Noordhoek JA, Timens W, Oosterhout AJM van, Postma DS. Abnormalities in Airway Epithelial Junction Formation in Chronic Obstructive Pulmonary Disease. American Journal of Respiratory and Critical Care Medicine. 2014 Jun 1. Located at: world. doi:10.1164/rccm.201311-1982LE

3. Raby KL, Michaeloudes C, Tonkin J, Chung KF, Bhavsar PK. Mechanisms of airway epithelial injury and abnormal repair in asthma and COPD. Front Immunol. 2023 Jul 13;14. doi:10.3389/fimmu.2023.1201658

4. Sayers I, John C, Chen J, Hall IP. Genetics of chronic respiratory disease. Nat Rev Genet. 2024 Aug;25(8):534–47. doi:10.1038/s41576-024-00695-0 PubMed PMID: 38448562.

5. Shrine N, Guyatt AL, Erzurumluoglu AM, Jackson VE, Hobbs BD, Melbourne CA, et al. New genetic signals for lung function highlight pathways and chronic obstructive pulmonary disease associations across multiple ancestries. Nat Genet. 2019 Mar;51(3):481–93. doi:10.1038/s41588-018-0321-7 PubMed PMID: 30804560; PubMed Central PMCID: PMC6397078.

6. Shrine N, Izquierdo AG, Chen J, Packer R, Hall RJ, Guyatt AL, et al. Multi-ancestry genome-wide association analyses improve resolution of genes and pathways influencing lung function and chronic obstructive pulmonary disease risk. Nat Genet. 2023 Mar;55(3):410–22. doi:10.1038/s41588-023-01314-0

7. Castaldi PJ, Cho MH, Litonjua AA, Bakke P, Gulsvik A, Lomas DA, et al. The Association of Genome-Wide Significant Spirometric Loci with Chronic Obstructive Pulmonary Disease Susceptibility. Am J Respir Cell Mol Biol. 2011 Dec;45(6):1147–53. doi:10.1165/rcmb.2011-0055OC

8. Cho MH, Boutaoui N, Klanderman BJ, Sylvia JS, Ziniti JP, Hersh CP, et al. Variants in FAM13A are associated with chronic obstructive pulmonary disease. Nat Genet. 2010 Mar;42(3):200–2. doi:10.1038/ng.535

9. Ehrhardt B, Roeder T, Krauss-Etschmann S. Drosophila melanogaster as an Alternative Model to Higher Organisms for In Vivo Lung Research. Int J Mol Sci. 2024 Sep 25;25(19):10324. doi:10.3390/ijms251910324 PubMed PMID: 39408654; PubMed Central PMCID: PMC11476989.

10. Ehrhardt B, El-Merhie N, Kovacevic D, Schramm J, Bossen J, Roeder T, et al. Airway remodeling: The Drosophila model permits a purely epithelial perspective. Front Allergy. 2022 Sep 15;3. doi:10.3389/falgy.2022.876673

11. Roeder T, Isermann K, Kabesch M. Drosophila in Asthma Research. Am J Respir Crit Care Med. 2009 Jun;179(11):979–83. doi:10.1164/rccm.200811-1777PP

12. Roeder T, Isermann K, Kallsen K, Uliczka K, Wagner C. A Drosophila asthma model - what the fly tells us about inflammatory diseases of the lung. Adv Exp Med Biol. 2012;710:37–47. doi:10.1007/978-1-4419-5638-5_5 PubMed PMID: 22127884.

13. Canales Coutiño B, Cornhill ZE, Couto A, Mack NA, Rusu AD, Nagarajan U, et al. A Genetic Analysis of Tumor Progression in Drosophila Identifies the Cohesin Complex as a Suppressor of Individual and Collective Cell Invasion. iScience. 2020 Jun 26;23(6):101237. doi:10.1016/j.isci.2020.101237

14. Couto A, Mack NA, Favia L, Georgiou M. An apicobasal gradient of Rac activity determines protrusion form and position. Nat Commun. 2017 May 19;8(1):1. doi:10.1038/ncomms15385

15. Georgiou M, Baum B. Polarity proteins and Rho GTPases cooperate to spatially organise epithelial actin-based protrusions. Journal of Cell Science. 2010 Apr 1;123(7):1089–98. doi:10.1242/jcs.060772

16. Canales Coutiño B, Szamek E, Markus Z, Georgiou M. Generation and live imaging of tumors with specific genotypes in the living fly pupa. STAR Protocols. 2021 Sep 17;2(3):100672. doi:10.1016/j.xpro.2021.100672

17. Schindelin J, Arganda-Carreras I, Frise E, Kaynig V, Longair M, Pietzsch T, et al. Fiji: an open-source platform for biological-image analysis. Nat Methods. 2012 Jun 28;9(7):676–82. doi:10.1038/nmeth.2019 PubMed PMID: 22743772; PubMed Central PMCID: PMC3855844.

18. Kim WJ, Lim JH, Lee JS, Lee SD, Kim JH, Oh YM. Comprehensive Analysis of Transcriptome Sequencing Data in the Lung Tissues of COPD Subjects. Int J Genomics. 2015;2015:206937. doi:10.1155/2015/206937 PubMed PMID: 25834810; PubMed Central PMCID: PMC4365374.

19. Brand AH, Perrimon N. Targeted gene expression as a means of altering cell fates and generating dominant phenotypes. Development. 1993 Jun 1;118(2):401–15. doi:10.1242/dev.118.2.401

20. Werder RB, Zhou X, Cho MH, Wilson AA. Breathing new life into the study of COPD with genes identified from genome-wide association studies. European Respiratory Review. 2024 May 29;33(172). doi:10.1183/16000617.0019-2024

21. Nishida K, Brune KA, Putcha N, Mandke P, O’Neal WK, Shade D, et al. Cigarette smoke disrupts monolayer integrity by altering epithelial cell-cell adhesion and cortical tension. American Journal of Physiology-Lung Cellular and Molecular Physiology. 2017 Sep;313(3):L581–91. doi:10.1152/ajplung.00074.2017

22. Shaykhiev R, Otaki F, Bonsu P, Dang DT, Teater M, Strulovici-Barel Y, et al. Cigarette smoking reprograms apical junctional complex molecular architecture in the human airway epithelium in vivo. Cell Mol Life Sci. 2011 Mar;68(5):877–92. doi:10.1007/s00018-010-0500-x PubMed PMID: 20820852; PubMed Central PMCID: PMC3838912.

23. Chen B, Brinkmann K, Chen Z, Pak CW, Liao Y, Shi S, et al. The WAVE Regulatory Complex Links Diverse Receptors to the Actin Cytoskeleton. Cell. 2014 Jan 16;156(1):195–207. doi:10.1016/j.cell.2013.11.048 PubMed PMID: 24439376.

24. Silva JM, Ezhkova E, Silva J, Heart S, Castillo M, Campos Y, et al. Cyfip1 Is a Putative Invasion Suppressor in Epithelial Cancers. Cell. 2009 Jun 12;137(6):1047–61. doi:10.1016/j.cell.2009.04.013 PubMed PMID: 19524508.

25. Costa PJ, Arndt KM. Synthetic Lethal Interactions Suggest a Role for the Saccharomyces cerevisiae Rtf1 Protein in Transcription Elongation. Genetics. 2000 Oct;156(2):535–47. doi:10.1093/genetics/156.2.535

26. Mueller CL, Jaehning JA. Ctr9, Rtf1, and Leo1 Are Components of the Paf1/RNA Polymerase II Complex. Molecular and Cellular Biology. 2002 Apr 1;22(7):1971–80. doi:10.1128/MCB.22.7.1971-1980.2002 PubMed PMID: 11884586.

27. Langenbacher AD, Lu F, Crisman L, Huang ZYS, Chapski DJ, Vondriska TM, et al. Rtf1 Transcriptionally Regulates Neonatal and Adult Cardiomyocyte Biology. Journal of Cardiovascular Development and Disease. 2023;10(5). doi:10.3390/jcdd10050221

28. Donaldson JG, Jackson CL. ARF family G proteins and their regulators: roles in membrane transport, development and disease. Nat Rev Mol Cell Biol. 2011 Jun;12(6):362–75. doi:10.1038/nrm3117

29. Mazelova J, Astuto-Gribble L, Inoue H, Tam BM, Schonteich E, Prekeris R, et al. Ciliary targeting motif VxPx directs assembly of a trafficking module through Arf4. The EMBO Journal. 2009 Feb 4;28(3):183–92. doi:10.1038/emboj.2008.267

30. Nakata K, Sugi Y, Narabayashi H, Kobayakawa T, Nakanishi Y, Tsuda M, et al. Commensal microbiota-induced microRNA modulates intestinal epithelial permeability through the small GTPase ARF4. Journal of Biological Chemistry. 2017 Sep 15;292(37):15426–33. doi:10.1074/jbc.M117.788596 PubMed PMID: 28760826.

31. Hsu SC, TerBush D, Abraham M, Guo W. The Exocyst Complex in Polarized Exocytosis. In: International Review of Cytology [Internet]. Academic Press; 2004 [cited 2024 Aug 28]. p. 243–65. Available from: https://www.sciencedirect.com/science/article/pii/S0074769604330068 doi:10.1016/S0074-7696(04)33006-8

32. Gervais L, Casanova J. In Vivo Coupling of Cell Elongation and Lumen Formation in a Single Cell. Current Biology. 2010;20(4):359–66. 10.1016/j.cub.2009.12.043

33. Jones TA, Nikolova LS, Schjelderup A, Metzstein MM. Exocyst-mediated membrane trafficking is required for branch outgrowth in Drosophila tracheal terminal cells. Developmental biology. 2014 Jun 6;390(1):41. doi:10.1016/j.ydbio.2014.02.021 PubMed PMID: 24607370.

34. Wingen A, Carrera P, Ekaterini Psathaki O, Voelzmann A, Paululat A, Hoch M. Debris buster is a *Drosophila* scavenger receptor essential for airway physiology. Developmental Biology. 2017 Oct 1;430(1):52–68. doi:10.1016/j.ydbio.2017.08.018

35. Terzikhan N, Xu H, Edris A, Bracke KR, Verhamme FM, Stricker BHC, et al. Epigenome-wide association study on diffusing capacity of the lung. ERJ Open Res. 2021 Mar 15;7(1):00567–2020. doi:10.1183/23120541.00567-2020 PubMed PMID: 33748261; PubMed Central PMCID: PMC7957297.

36. O’Donohue MF, Choesmel V, Faubladier M, Fichant G, Gleizes PE. Functional dichotomy of ribosomal proteins during the synthesis of mammalian 40S ribosomal subunits. J Cell Biol. 2010 Sep 6;190(5):853–66. doi:10.1083/jcb.201005117 PubMed PMID: 20819938; PubMed Central PMCID: PMC2935573.

37. Wang W, Nag S, Zhang X, Wang MH, Wang H, Zhou J, et al. Ribosomal Proteins and Human Diseases: Pathogenesis, Molecular Mechanisms, and Therapeutic Implications. Medicinal Research Reviews. 2015;35(2):225–85. 10.1002/med.21327

38. Baker NE. Mechanisms of cell competition emerging from Drosophila studies. Current Opinion in Cell Biology. 2017;48:40–6. 10.1016/j.ceb.2017.05.002

39. Portelli MA, Rakkar K, Hu S, Guo Y, Adcock IM, Sayers I. Translational Analysis of Moderate to Severe Asthma GWAS Signals Into Candidate Causal Genes and Their Functional, Tissue-Dependent and Disease-Related Associations. Frontiers in Allergy [Internet]. 2021 [cited 2024 Jan 5];2. Available from: https://www.frontiersin.org/articles/10.3389/falgy.2021.738741

40. Kahvejian A, Svitkin YV, Sukarieh R, M’Boutchou MN, Sonenberg N. Mammalian poly(A)-binding protein is a eukaryotic translation initiation factor, which acts via multiple mechanisms. Genes Dev. 2005 Jan 1;19(1):104–13. doi:10.1101/gad.1262905 PubMed PMID: 15630022; PubMed Central PMCID: PMC540229.

41. Mangus DA, Evans MC, Jacobson A. Poly(A)-binding proteins: multifunctional scaffolds for the post-transcriptional control of gene expression. Genome Biol. 2003 Jul 1;4(7):223. doi:10.1186/gb-2003-4-7-223

42. Kamiyama T, Sun W, Tani N, Nakamura A, Niwa R. Poly(A) Binding Protein Is Required for Nuclear Localization of the Ecdysteroidogenic Transcription Factor Molting Defective in the Prothoracic Gland of Drosophila melanogaster. Frontiers in Genetics. 2020;11. doi:10.3389/fgene.2020.00636

43. Peng S, Fang J, Mo W, Hu G, Wu S. Identifying cross-tissue molecular targets of lung function by multi-omics integration analysis from DNA methylation and gene expression of diverse human tissues. BMC Genomics. 2025 Mar 24;26(1):289. doi:10.1186/s12864-025-11476-2

44. Maloy A, Walter S, Saferali A, Mascilli A, Natri H, Berman A, et al. PABPC4: A Novel COPD Susceptibility Gene Regulating Mitochondrial Function and Emphysema Pathogenesis Targeted by a First-in-Class Therapeutic Agent. Am J Respir Crit Care Med. 2025 May 1;211(Supplement_1):A7334. doi:10.1164/ajrccm.2025.211.Abstracts.A7334

