## Supplementary Methods, Figures, and Legends for "Lung function candidate genes in *Drosophila melanogaster*"

**Author information**

^1^School of Life Sciences, University of Nottingham, Nottingham, UK.

^2^ Department of Genetics, Microbiology and Statistics, Universitat de Barcelona (UB).

^3^ Institute of Biomedicine of the University of Barcelona (IBUB), Barcelona, Spain.

^4^Department of Thoracic Medicine, National Heart and Lung Institute, Imperial College School of Medicine, London, UK.

^5^Centre for Respiratory Research, NIHR Nottingham Biomedical Research Centre, School of Medicine, Biodiscovery Institute, University of Nottingham, Nottingham, UK.

*contributed equally.

**Supplementary Methods**

Online methods

**Candidate gene prioritisation**

At the time of analysis, the largest GWAS meta-analysis of lung function was by Shrine et al. (2019) which assembled genotype data from UK Biobank (N= 321,047) and the SpiroMeta consortium (n= 79,055) comprising over 400,102 individuals of European ancestry (1). In this study, 19,819,130 SNPs (genotyped and imputed) were meta-analysed and a total of 279 signals (139 new and 140 previously reported) were significantly associated with four quantitative lung function traits: FEV_1_, FVC, FEV_1_/FVC and PEF.

To identify causal variants in trait-associated loci and interpret their biological mechanisms, Shrine and colleagues applied a Bayesian method (2) of fine-mapping SNPs, and utilised gene and protein expression data. From these approaches 107 putative causal genes were identified, corresponding to 70 (30 novel, 40 previous) independent signals (out of a total 279) for further study.

In the current study we completed variant to gene mapping for the 70 signals that had mapped to candidate causal genes with the greatest confidence utilising publically available datsets (The Genotype Tissue Expression Consortium, GTEx v8; Open Target Genetics, OTG release 20.2; Ensembl; LD Matrix 5.0; HaploReg V4.1), and unique (Unbiased biomarkers for the prediction of respiratory disease outcomes, U-BIOPRED; bulk RNA-sequencing) datasets. We retrieved SNP-specific annotations including variant 2 gene (V2G) association scores, tissue-specific eQTLs and sQTLs, VEP and epigenomic information i.e., DNAse hypersensitivity sites (DHSS) and promoter high throughput chromosome conformation capture (PCHI-C) interactions. The identification of additional respiratory relevant traits associated with signals was achieved through a phenome-wide association study (PheWAS) analysis using the UK Biobank resource. These analyses provided a candidate causal gene list for further investigation.

To evaluate the human candidate gene list to orthologue associations, we consulted publicly accessible databases (The DRSC Integrative Ortholog Prediction Tool, DIOPT v8; Drosophila Gene Expression Tool, DGET 1.0.2; and FlyBase, FB2020_06 release).

**Immunolabelling**

Dissected nota were fixed in 4% PFA at room temperature (RT) for 20 minutes, rinsed three times in 0.1% PBS-Triton X (PBS-T), then transferred to a glass well containing blocking solution (0.2% BSA and 5% NGS in 0.1% PBST) at RT for an hour. PBS-T was subsequently replaced with primary antibody solution and samples stored overnight within a humidified chamber at 4°C. As previous, samples were rinsed three times with PBS-T before being incubated with secondary antibody for an hour at RT. Following this, three further washes with PBS-T took place. Samples were mounted individually on a microscope slide using GeneTex FluoroGel (#GTX28214) and Sigma mounting media (#F4680) with the apical (dorsal) domain facing upwards. A coverslip was overlayed, and samples left to set overnight at 4°C prior to imaging.

**Collection, fixation and immunolabeling of whole embryos**

To obtain *Drosophila* embryos, we set up fly crosses with 10 UAS-RNAi males and 25 btl-Gal4 virgin females of interest. Crosses were placed in an egg laying cage with a yeast-enriched agar plate at the base. The egg-collecting cage was incubated overnight at 25°C and 29°C when an increased activity of GAL4 was required. On collection, the leftover yeast was removed with the spatula and the plate filled with bleach for two minutes to dechorionate embryos. During this time, embryos were carefully brushed onto a mesh collecting basket and rinsed with Triton solution. Excess liquid was removed from embryos by placing the collection basket on dry tissue paper. Embryos were subsequently transferred to a vial containing fixative solution: 4% formaldehyde, PBS (0,1 M NaCl 10mM phosphate buffer, pH 7,4) and heptane [1:1] for 20 minutes at RT. The solution was shaken to allow formaldehyde to partition into a heptane phase. Embryos were retrieved from the interface between the two phases with a Pasteur pipette, washed 2-3 times with methanol and left overnight at 4°C. Following this, embryos were rinsed three times with PBT (PBS 1% con 0,1% Tween 20), then resuspend in PBT and PBT-BSA (1% Bovine Serum Albumin) for 30 minutes respectively. These embryos were placed at 4°C overnight with primary antibody [A2A12, mouse anti-GASP] diluted as shown in the Appendix 7.1. Post incubation, embryos underwent three aforementioned washes with PBT and were stained with secondary antibody [1:500, mouse IgM (anti-2A12)] diluted in PBT+BSA for two hours at RT. As previous, embryos were rinsed and washed twice for 20 min with PBT.

For colour detection, a 1:200 (A) avidin and (B) Biotinylated Horseradish Peroxidase H solution was prepared using the Vectastain® ABC kit (Vector laboratories) according to the supplier’s protocol. The solution was applied to embryos and incubated for 30 minutes at RT. After three rinses and a further PBT wash step for 10 minutes, 500μl of 3’3 diaminobenzidine (DAB) was added. This reaction monitored until a brown precipitate formed (typically 3 minutes) and deactivated with three PBT rinses. Prior to mounting, stained embryos were stored in 70% glycerol.

**Developmental Viability Assays**

Viability assays were carried out to assess the fitness of specific genotypes by observing overall lethality at each stage of development. Crosses set up between UAS-RNAi line and btl-Gal4 genotypes were stored at 25°C overnight and flipped into a new vial a day later. Emptied vials were monitored over a 20-day period to determine whether embryos would progress to adulthood or die prematurely. This procedure was repeated to obtain three experimental replicates.

**Quantification of screen phenotypes**

To perform quantitative analysis, image files (czi or TIFF file format) were loaded into Fiji ImageJ software. Separately, a spreadsheet was set up in Excel using formulae to automate the calculations below.

Clonal area:

To measure clonal area, the polygon tool was used to trace the outline of all visible clones in a z-projected image (Image>Stacks>Z-project, projection intensity=maximum). Between each clone, measurements (Analyse>Measure) were saved before adding all values together.

% Clonal area = "Total clonal area" /"Total area (354.25 x 354.25)" "x100"

Delaminated cells and dividing cells:

To approximate the number of clonal cells in an image, a z-projected image of the apical slice/s was made (Image>Stacks>Z-project, projection intensity=maximum). A 35um2 box was drawn (Edit>Selection>Specify), positioned over image, and the number of cells counted in the enclosed area using the multipoint tool.

Total number of clonal cells = "Total clonal area" /"(35 x 35)" x number of cells in 35um^2^

To quantify the number delaminated or dividing cells, the multipoint tool is used to count cells throughout the entire z-stack, saved and input into the following formula:

% Delaminated cells="Number of delaminated cells (whole interface)" /"Total number of clonal cells (previous)" "x100"

**Quantification of junctional fluorescence**

To quantify fluorescent intensity at the junctions of isolated immunostained nota in distinct regions i.e. KD clones *versus* wildtype areas requires image preprocessing, mask creation and line profiling which was carried out by the following devised method:

Image files (czi file format) were loaded into Fiji ImageJ software. An editable copy of the full image was generated (Image>Duplicate>Duplicate hyperstack). Z-slices were maximally projected (Image>Stacks>Z-project, projection intensity=maximum) and a fixed subset of slices specified to eliminate noise from the basal region. The composite image was separated into constitutive GFP and cadherin channels (Image>Color>Split channels). As a preprocessing step (Process>Filter), Gaussian Blur (sigma value =1.5) was applied to the GFP-positive image. A mask (Image>Adjust>Threshold) from the blurred image was generated with the automated thresholding program (‘Triangle’ and ‘Dark background’) to produce an 8-bit binary image. The mask was further refined (Process>Binary>Fill holes, ‘Open’) and pixel values checked on the outside of the clone (black=0) and inside the clone (white=255). To turn the mask into a multiplier, values are subtracted from the clonal region (Process>Math>Subtract value=254), resulting in a clonal region pixel value of 1. A further visualisation step was taken (Adjust>Brightness/Contrast, shift ‘Maximum’ slider to the left).

To match the GFP-positive clonal region to an equivalent area in the cadherin channel, the mask window was multiplied by the template cadherin image (Process>Image calculator>Multiply). The output punches a hole in the red channel, eliminating WT areas (multiplied by zero), leaving the clonal area outline on the cadherin image for analysis. For improved visualisation and figure preparation, images were converted to grayscale (Image>Lookup Tables>Grays). To minimise selection bias, all cells were numbered using the multi-point tool. Numbers were then randomised using the RANDDBETWEEN formula in Excel. To measure fluorescent intensity at a given junction/s, the image was magnified to 300% and the segmented line tool used to trace the length of a junction. The mean gray value, SD, minimum/maximum intensity values were recorded (Analyse>Measure). Importantly, to account for the morphological influence of the neighbouring WT cells, cells on the boundary between of clonal and WT tissue were not included in analysis.

1.	Shrine N, Guyatt AL, Erzurumluoglu AM, Jackson VE, Hobbs BD, Melbourne CA, et al. New genetic signals for lung function highlight pathways and chronic obstructive pulmonary disease associations across multiple ancestries. Nat Genet. 2019 Mar;51(3):481–93. doi:10.1038/s41588-018-0321-7 PubMed PMID: 30804560; PubMed Central PMCID: PMC6397078.

2.	Wakefield J. Reporting and interpretation in genome-wide association studies. International Journal of Epidemiology. 2008 Jun 1;37(3):641–53. doi:10.1093/ije/dym257

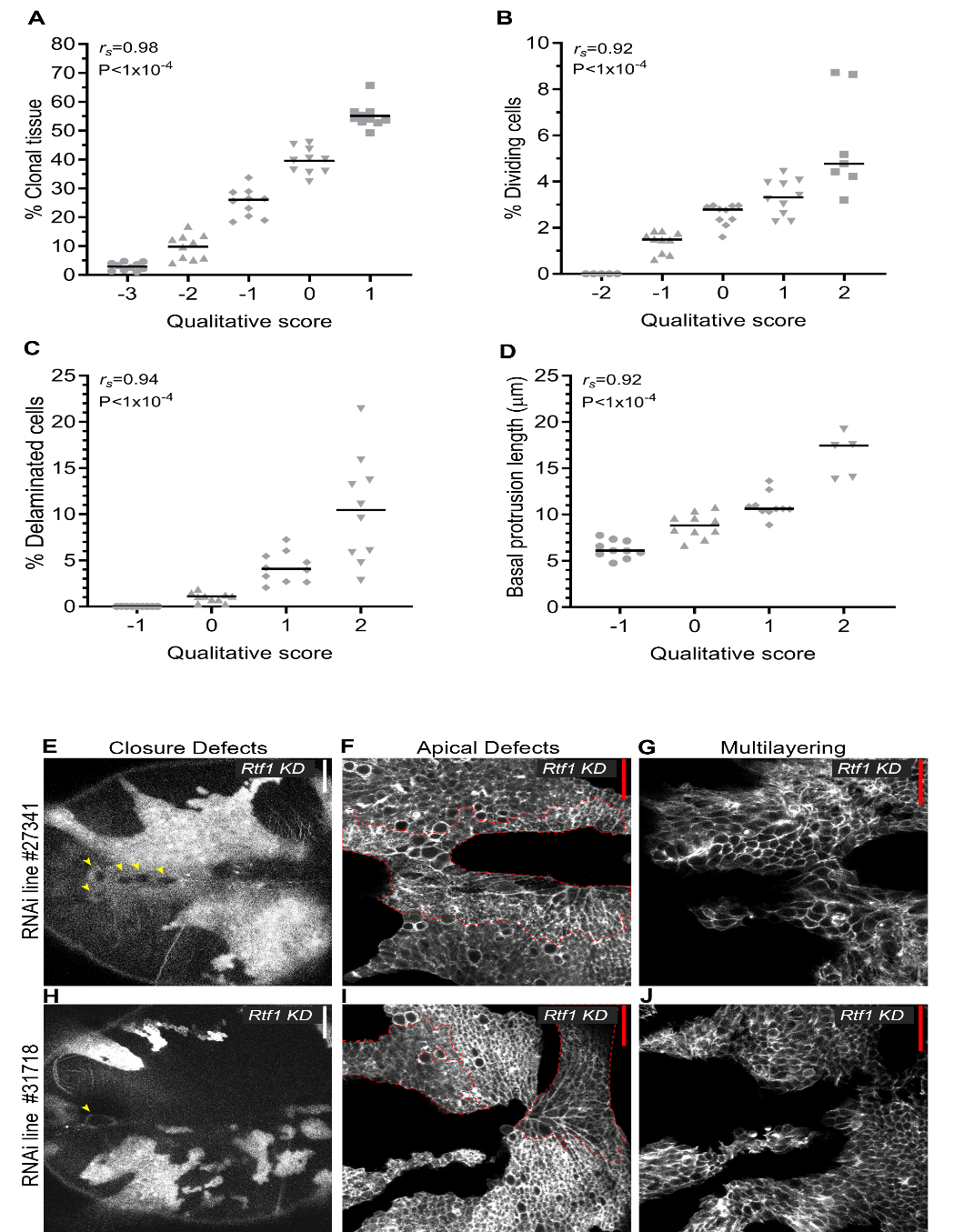

Supplementary Figures

**Figure S1** Quality control. (A-D) Graphs comparing semi-quantitative scores given to individual animals against the quantitative measure for a range of phenotypes, including: (A) percentage clonal tissue; (B) percentage dividing cells; (C) percentage invading cells and (D) basal protrusion length (n=10 animals for each category of qualitative score with three exceptions). (E-J) Image panels comparing the knockdown phenotypes of two independent UAS-RNAi lines (#27341, #31718) targeting gene Rtf1 in categories closure defects, apical defects and multilayering. UAS-RNAi lines were obtained from three different near genome wide libraries: VDRC in Austria, NIG in Japan, and Bloomington in the USA. White scale bar (E, H) = 50μm; red scale bar (F-G, I-J) = 30μm; yellow arrows (E, H) indicate position of closure defect, red outline (F, I) encloses a defective apical tissue within a clone.

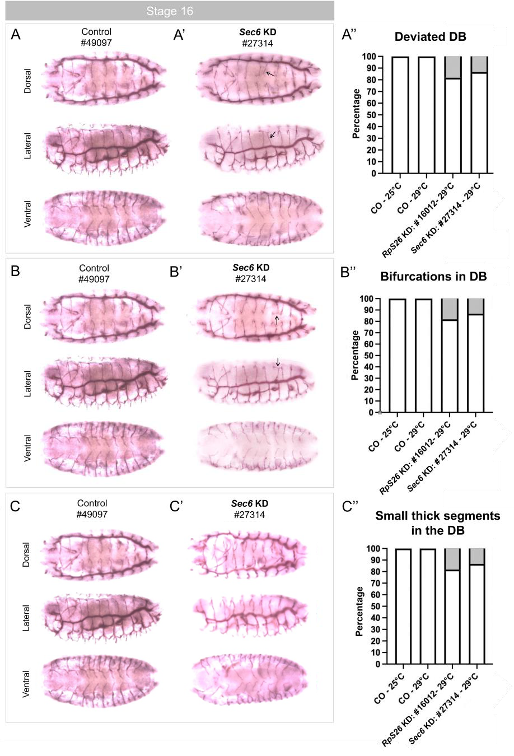

**Figure S2** Characterisation of tracheal phenotypes in knockdown embryos. Non-significant numbers of knockdown induced tracheal defects present in embryonic stages development. (A-C) Representative control embryos and (A’-C’) Sec6 knockdown (RNAi #27314) embryos visualised at stage 16 of embryonic development using a lumen-specific mAb2A12 antibody. Following Sec6 and RpS26 knockdown, tracheal phenotypes were observed in the dorsal branches (DB) including (A’; black arrows at DB6) deviated dorsal branches; (B’; black arrows at DB7) bifurcation of a terminal branches; and (C’; black arrows) small thicker segments near the dorsal trunk. Anterior view: left, dorsal: up. (A’’-C’’) Percentage of total RNAi lines embryos displaying tracheal phenotypes versus controls. Images were obtained using a Leica microscope. 10 ≤ n ≤ 20. *P*<0.05.

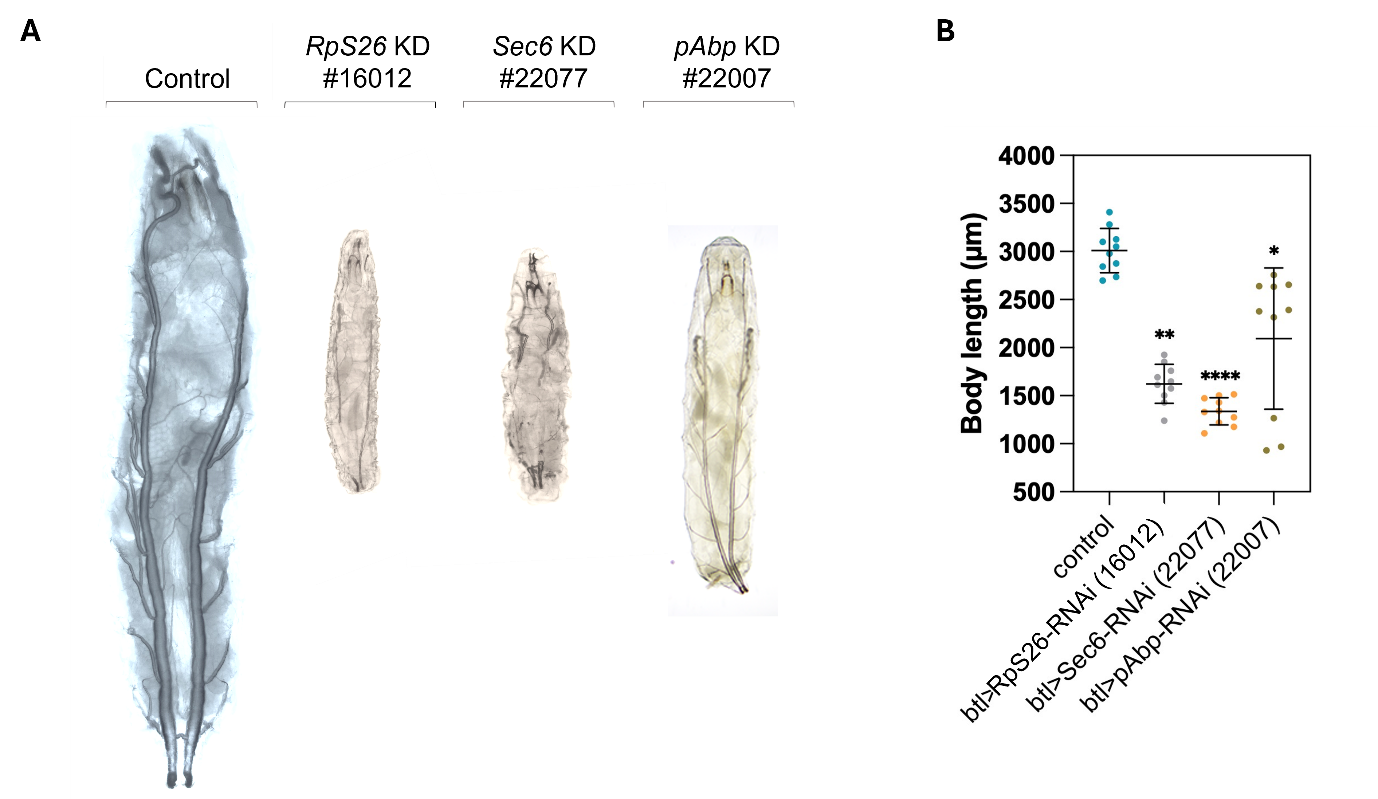

**Figure S3** Knockdown of *RpS26*, *Sec6* and *pAbp* result in tracheal alterations coupled with a significant reduction in larval body size. (A) Representative images *RpS26*, Sec6 and pAbp L3 knockdown larvae displaying a combination of phenotypes including significant tracheal defects including a loss of continuity in the dorsal trunks, thinner airways and a smaller body size relative to control larvae. (B) Quantifications of larval body length. Images were obtained using a Leica microscope. n (number of embryos studied) =10. Mann-Whitney U-test **P*<0.01; ***P*<0.001; *****P*<0.0001.
